# Nanopore Direct RNA Sequencing Reveals Virus-Induced Changes in the Transcriptional Landscape in Human Bronchial Epithelial Cells

**DOI:** 10.1101/2024.06.26.600852

**Authors:** Dongyu Wang, J. Leland Booth, Wenxin Wu, Nicholas Kiger, Matthew Lettow, Averi Bates, Chongle Pan, Jordan Metcalf, Susan J. Schroeder

## Abstract

Direct RNA nanopore sequencing data reveals changes in gene expression, polyadenylation, splicing, m6A methylation, and pseudouridylation in response to influenza virus exposure of primary human bronchial epithelial cells. This study focuses on the epitranscriptomic profile of genes in the host immune response. In addition to polyadenylated noncoding RNA, we purified and sequenced nonpolyadenylated noncoding RNA and observed changes in expression, N6-methyl-adenosine (m6A), and pseudouridylation (Ψ) in these novel RNA. Two recently discovered lincRNA with roles in immune response, *Chaserr* and *LEADR*, are predicted in the analysis of the sequencing data to be highly methylated in response to influenza exposure. Several H/ACA type snoRNAs that guide pseudouridylation are decreased in expression in response to influenza, and there is a corresponding predicted decrease in the pseudouridylation of two novel lncRNA. Thus, novel epitranscriptomic changes revealed by direct RNA sequencing with nanopore technology provide unique insights into the host epitranscriptomic changes in epithelial gene networks that respond to influenza virus exposure.

## Introduction

Seasonal influenza and pandemic influenza cause significant morbidity and mortality. The Centers for Disease Control estimates that 9 million – 41 million illnesses, 140,000 – 710,000 hospitalizations and 12,000 – 52,000 deaths annually between 2010 and 2020 were due to influenza ^1^. Viruses are master manipulators of host cellular resources. Changes in ribonucleic acid (RNA) provide the genetic instructions for a host cell to respond to a virus infection or exposure. The changes in modified nucleotides can alter splicing and the way that the RNA folds and interacts with proteins and thus fine-tune host gene regulation responses to viruses but may also be modulated by viruses to promote infection. The epitranscriptome includes all the changes in modified nucleotides in response to cell stresses and developmental gene regulation. This manuscript presents a study of the transcriptomic and epitranscriptomic changes in response to influenza virus exposure in primary human bronchial epithelial cells (HBEC) using direct RNA sequencing from Oxford Nanopore Technology and reveals changes in RNA abundance, nucleotide modifications, splicing isoforms, and polyA tail length in response to influenza. We identify changes in RNA methylations in both coding and noncoding RNA. Direct RNA nanopore sequencing (DRS) is a single molecule, long-read sequencing technology that measures nucleotide modifications, splicing isoforms, polyA tail length, and differential expression in a single assay. This new method, therefore, demonstrates the changes in viral and host RNA that occur during IAV exposure and the host immune response at higher resolution than standard methods, enhancing our understanding of these processes.

RNA adenosine methylation is an important regulatory mechanism of the transcriptional response to viral infection or exposure. N6-methyl-adenosine (m6A) is the most common modified RNA nucleotide, and the levels of m6A change globally in response to many types of viral infections ^2–5^. m6A modifications regulate splicing^6^, translational efficiency, RNA folding and tertiary interactions, RNA-protein interactions, and RNA cellular half-life in human transcriptomes ^7–9^. m6A modulates host response through several different gene networks. IAV induces a strong interferon (IFN) immune response through Interferon Regulatory Factor (IRF) 7 activation in human lung bronchial and alveolar epithelial cell models ^10^. Adenosine methylation negatively regulates the type I interferon response through m6A modification of *IFNβ* mRNA in many kinds of viral infection, including influenza infection ^11^. The loss of m6A methylation on *IFNβ* mRNA increases mRNA stability and thus causes a more sustained IFN-β production and a more robust antiviral response. In Herpes Simplex Virus 1 infection, m6A methylation increases mRNA half-lives of the immune response genes *IFIH1*, *TNFIP3*, *IFIT1* and reduces the half-life of *IFIT2* mRNA ^12^. In Kaposi’s sarcoma-associated herpesvirus infection, methylation of IL-6 transcripts in the 3’UTR recruit the YTHDC2 reader, preventing viral SOX-induced mRNA degradation ^3^, again increasing the half-life of this transcript. In adenovirus viral RNA transcripts, m6A methylation regulates the splicing efficiency of late-stage genes ^13^. In developing attenuated Respiratory Syncytial Virus (RSV) vaccines, it was discovered that RSV viral RNA that lacks m6A binds RIG-I with greater affinity, induces higher RIG-I expression and ubiquitination, increases IRF3 phosphorylation, increases RIG-I IFN signaling, and enables a higher IFN response ^14^. m6A methylation in the 3’ UTR region can either stabilize or destabilize the cellular lifetime of the RNA transcript. For example, m6A in the 3’UTR and coding regions of *STAT1* increases the cellular stability of *STAT1*, a key master regulator of macrophage differentiation ^15^. In contrast, m6A methylation of the 3’UTR in the *PTX3* gene increases the degradation of the RNA transcript and results in the repression of M2 activation ^16^. Previous studies have shown that m6A changes globally in response to virus infection ^17,18^. Our data in HBEC cells uses direct RNA nanopore sequencing to identify m6A changes in the context of full-length, single-molecule transcript reads revealing epitranscriptomic changes in the immune response gene network.

Traditional studies of host and virus RNA methylations in influenza infections have used mass spectrometry or variations of Illumina sequencing with m6A specific antibodies to pull down modified RNA fragments ^9,11,18–22^. Direct RNA sequencing with nanopore technology offers a new window to study RNA methylations directly in long sequencing reads. Nanopore sequencing technology uses a motor protein in a pore in a synthetic lipid bilayer to transfer a single molecule of DNA or RNA across a membrane. The voltage in picoamperes is measured as a function of time and sampled at 4 KHz. The voltage varies depending on the sequence of 5-6 nucleotides that are located in the channel of the pore and the corresponding pH difference on each side of the membrane. Modified nucleotides such as m6A alter the pK_a_ of the adenine base and correspondingly change the signal in the nanopore data. Modified nucleotides cause changes in the signal intensity, duration, and frequency of base calling errors ^23,24^. m6Anet uses a multiple instance learning framework to analyze the nanopore signals to predict the probability and abundance of an m6A modified nucleotide at a specific nucleotide position in a transcript^25^. m6Anet currently outperforms other analysis methods in accuracy ^26^, although research and the development of more accurate tools is ongoing.

Nanospa software identifies both m6A and pseudouridine in DRS nanopore data^27^. Some prior studies have focused on pseudouridine in interferon responses^28,29^; stress responses ^30^; and in influenza genomic viral RNAs^20,31^. However, much less is known about pseudouridine in host mRNA or lncRNA in IAV infections or exposure. Nanopore sequencing is currently being used to directly sequence and monitor influenza RNA. Oxford Nanopore Technology has developed ARTIC protocols to identify novel, emergent RNA viruses and new sequences of influenza viral RNA ^32,33^. The variations in m6A in influenza viral mRNA were first observed using RNA digestion and high pressure liquid chromatography ^34^. m6A methylation of some influenza viral transcripts enhances viral replication ^20^. More recently, nanopore sequencing of the influenza coding genome used specific adapter sequences to detect the viral RNA in cultured cells ^35^. Since the first application of Direct RNA Sequencing (DRS) to study influenza, significant improvements in basecalling software Guppy and Dorado ^36^ and identification of m6A using machine learning programs such as m6Anet have advanced nanopore DRS. Both host and viral RNA change methylations in response to virus infection ^18^. This nanopore DRS study focuses on the host immune response to viral exposure and the local and global changes in m6A methylation in coding and noncoding RNA.

## Materials and Methods

### Cell culture

Human bronchial epithelial cells (HBEC) were collected from three clinical samples donated by healthy, adult non-smokers of diverse ages, genders, and races. All samples were collected following protocols that were approved by the Institutional Review Board of the University of Oklahoma Health Sciences Center (IRB # 2197). HBEC were obtained by bronchoscopy with bronchial brushing. Three or four separate third or fourth order bronchi were brushed using a sterile cytology brush and the cells were rinsed from the brush into 10 ml sterile saline until 5×10^6^ to 1×10^7^ cells total were collected as determined by hemocytometer counts for total viable cells as assessed by trypan blue exclusion. The cells for isolation were resuspended to 5×10^5^ cells/ml in complete Bronchial Epithelial Cell Growth Medium (BEGM; Lonza Group Ltd.); were seeded into collagen coated tissue culture plates (Bio-Coat, BD Biosciences) at a density of 1×10^5^ cells/cm^2^ and were propagated in an incubator at 37°C in 5% CO_2_. After 24 h the cells are washed with HBSS to remove non-adherent cells and fresh complete BEGM is added. When the cultures are near confluence (7-10 days), the monolayers are lifted with 1x Accutase solution and are subcultured at a 1:5 dilution. After each passage the cells grow to confluence within 4 to 5 days, and when the cultures are split, freezer stocks are prepared in 80% BEGM + 10% fetal bovine serum + 10% DMSO and are stored in liquid nitrogen vapor at -190°C.

### HBEC exposure, Virus stock preparation

The HBEC were infected with influenza as previously described^10,37^. H1N1 influenza A virus, A/PR/34/8 (PR8), was passaged in Madin-Darby canine kidney (MDCK) cells. Virus was propagated in MDCK cells in DMEM/F12 with ITS+ (BD Biosciences, Franklin Lakes, NJ) and trypsin, harvested at 72 h post-exposure and titered by plaque assay in MDCK cells. The virus was stored in aliquots at −80 °C. No detectable endotoxin was observed in the final viral preparations, as determined by limulus amebocyte lysate assay (Cambrex, Walkersville, MD) with a lower limit of detection at 0.1EU/ml or approximately 20 pg/ml LPS. HBEC cultures were exposed to IAV at a multiplicity of infection (MOI) of 1. Total cellular RNA was collected 24 hours post-exposure.

### RNA preparation

The total RNA from cells was extracted with Trizol using Zymo columns with modifications described below and then assessed for quantity on a Nanodrop spectrometer and quality on an Agilent tape station. Sample N24 was further ethanol precipitated with sodium acetate (0.1 volume of sodium acetate pH 5.2 and 2 volumes of ethanol added to the sample with incubation at -80 °C, followed by centrifugation at 4 °C and a wash with 70% v/v ethanol at 4 °C) to remove traces of Trizol reagent. 4-50 μg of total cellular RNA was polyA purified using a Lexogen polyA RNA selection kit with elution into 12 μL volume. The quality of RNA and rRNA depletion was assessed using an Agilent tape station. The RNA from step 9 of the polyA purification protocol was further purified using a Lexogen Ribocop kit in order to collect nonpolyadenylated RNA enriched in ncRNA and depleted of ribosomal RNA. The resulting RNA was then *in vitro* polyadenylated using New England Biolabs polyA polymerase from *E. coli* (product # M0276S) with incubation at 37 °C for 90 minutes and again purified using the polyA selection kit. Oxford Nanopore kit ONT RNA-SQK002 protocols including the reverse transcription step were used to prepare libraries for nanopore sequencing with 180 ng - 2μg input RNA. Oxford Nanopore R9.4 minion flow cells and the Oxford MinION platform (MinKnow 22.12.7) were used for sequencing.

### Direct RNA nanopore sequencing data analysis

Oxford nanopore base calling software, Guppy ^38^ within Dorado version 0.3.2 ^36^, was used for high quality basecalling with a minimum quality score of 7 for passing reads. Minimap 2 algorithm ^39^ was used to align RNA sequences to the human transcriptome, GENCODE v45 ^40^. FLAIR v1.4 (Full Length Alternative Isoform analysis of RNA) ^41^ was used for analysis of alternative splicing. m6Anet software ^25^ was used for analysis of m6A methylation to determine sites in a sequence that are likely to be m6A methylated. m6Anet requires a minimum of 30 copies for a transcript to be analyzed. m6Anet output includes the transcript ID, nucleotide number, probability of m6A, and percent of transcripts with m6A at that nucleotide site. Only sites with an m6A probability equal to or greater than 0.90 were included in further analysis. NanoSPA software was used to identify m6A and pseudouridine modifications^42^. Both m6Anet and NanoSPA were trained on DRS data from human RNA samples. Although these tools can sometimes overpredict sites, the probability level cutoffs were conservative in order to identify well-predicted sites. Further validation of predictions from m6Anet and NanoSPA tools uses using mass spectrometry and synthetic libraries and is a continuing, ongoing area of research^43^. Differential expression was determined using DESeq2 ^44^. The data has been deposited to the European Nucleic Acid Archive PRJNA1134237.

### Quantitative Reverse Transcription PCR Analysis

In order to validate differential gene expression in the ncRNA, quantitative Reverse Transcription PCR was done for two snoRNA in total cellular RNA. One microgram of total cellular RNA was used for cDNA synthesis. In order to remove any traces of genomic DNA, the RNA was treated with TurboDNAse for 1 hour at 37 °C, inactivated, centrifuged, and stored at -20 °C following manufacturer’s protocol. 300 ng of RNA was annealed with 500 ng of random primers from Promega for 5 minutes at 70 °C. 300 U of M-MLV reverse transcriptase (Promega M170) was added with buffer and NTPs, as per manufacturer’s protocols, and incubated at 25 °C for 15 min, 42 °C for 1 h, 95 °C for 2 min, and then 4 °C. Approximately 3 ng of cDNA and 0.5 μM primers were mixed with 2X SYBR Green supermix in a 25 μL reaction. A Biorad CFX Connect Real Time System and the Biorad Sso Advanced SYBR Green protocols were used for data collection and analysis. The PCR program was 95 °C for 3 min, 45 cycles of 95 °C for 10s and 50 °C for 30 s, and then melting from 65 °C to 95 °C in 0.5 °C increments. Data was analyzed using Biorad CFX Manager software and the ΔΔCt method with β-actin as the reference gene. The primers for β-actin, snoRA23, and snoRA33 are listed in Supporting Information Table 2.

## Results and Discussion

### m6A methylation, polyA tail length, and alternative splicing of mRNA transcripts in HBECs are altered by influenza exposure

In our DRS studies of influenza exposure in HBEC cells, we identified global changes in gene expression, alternative splicing, m6A methylation, and polyA tail length between influenza-exposed and mock-exposed HBECs. We detected 1,473 and 2,729 transcripts with 2 or more isoforms and 502 and 395 transcripts with predicted m6A methylation in exposed and unexposed HBEC cells, respectively. DRS analysis of the 3 paired data sets shows high quality RNA sequencing for polyA transcripts. Table 1 shows the overview of the results of DRS analysis in polyA and nonpolyadenylated transcripts. The figures showing high accuracy basecalling at >95% are shown in Figure S1. Figures showing the accuracy of mapping reads in plots of normalized read identity versus read length are also shown in Figure S1.

**Table 1.** Direct RNA Sequencing Analysis of Transcripts in Human Bronchial Epithelial Cells Exposed to IAV. The p-value was adjusted as p-adjust by the method of Benjamini-Hochberg^70^. The cut off for p-adjust is 0.05. The cutoff of for log2 fold change in differentially expressed genes is 1.2. The lowest probability cutoff for m6A methylation in m6Anet is 0.90.

| sample | reads mapped | reads with m6A | reads > 1 alternative splicing event | Differentially expressed reads | reads with differential polyA tail |
| --- | --- | --- | --- | --- | --- |
| polyA transcripts |  |  |  |  |  |
| Donor 1 uninfected | 1448955 | 4129 | 10626 | 11527 | 3037 |
| Donor 1 Infected | 1548820 | 6657 | 9383 |  |  |
| Donor 2 uninfected | 669296 | 1695 | 5797 |  |  |
| Donor 2 Infected | 442279 | 4205 | 5434 |  |  |
| Donor 3 uninfected | 200368 | 236 | 2389 |  |  |
| Donor 3 Infected | 282294 | 196 | 2711 |  |  |
| Nonpolyadenylated transcripts |  |  |  |  |  |
| Donor 1 uninfected | 8909 | 0 | 0 | 44 | Not applicable |
| Donor 1 Infected | 748 | 0 | 0 |  |  |
| Donor 2 uninfected | 112016 | 1 | 0 |  |  |
| Donor 2 Infected | 2224 | 0 | 0 |  |  |

**Table 2A:**
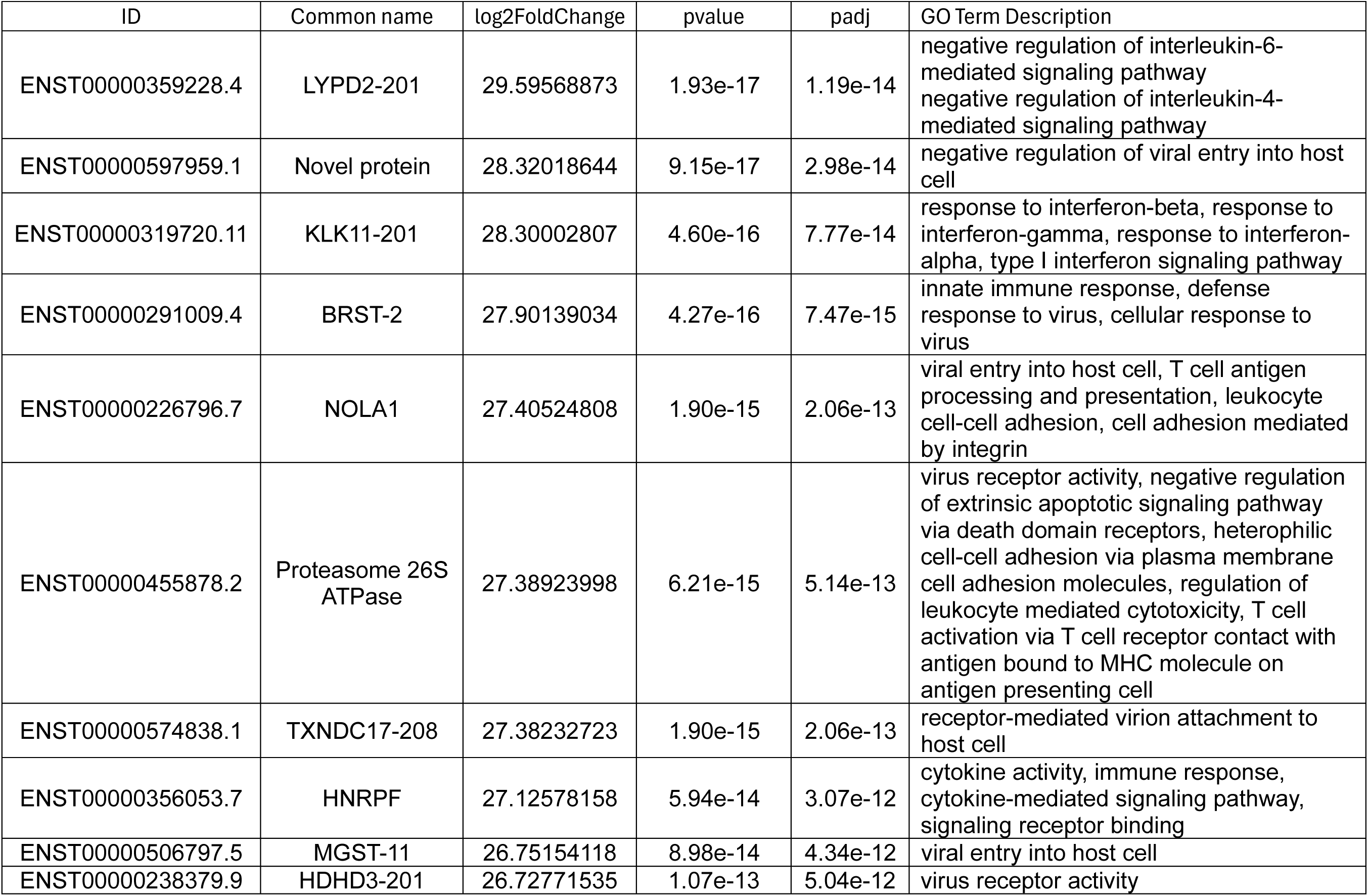
Transcripts Up-Regulated in Response to IAV Exposure and Related to Immune Response and Virus Entry.

**Table 2B:**
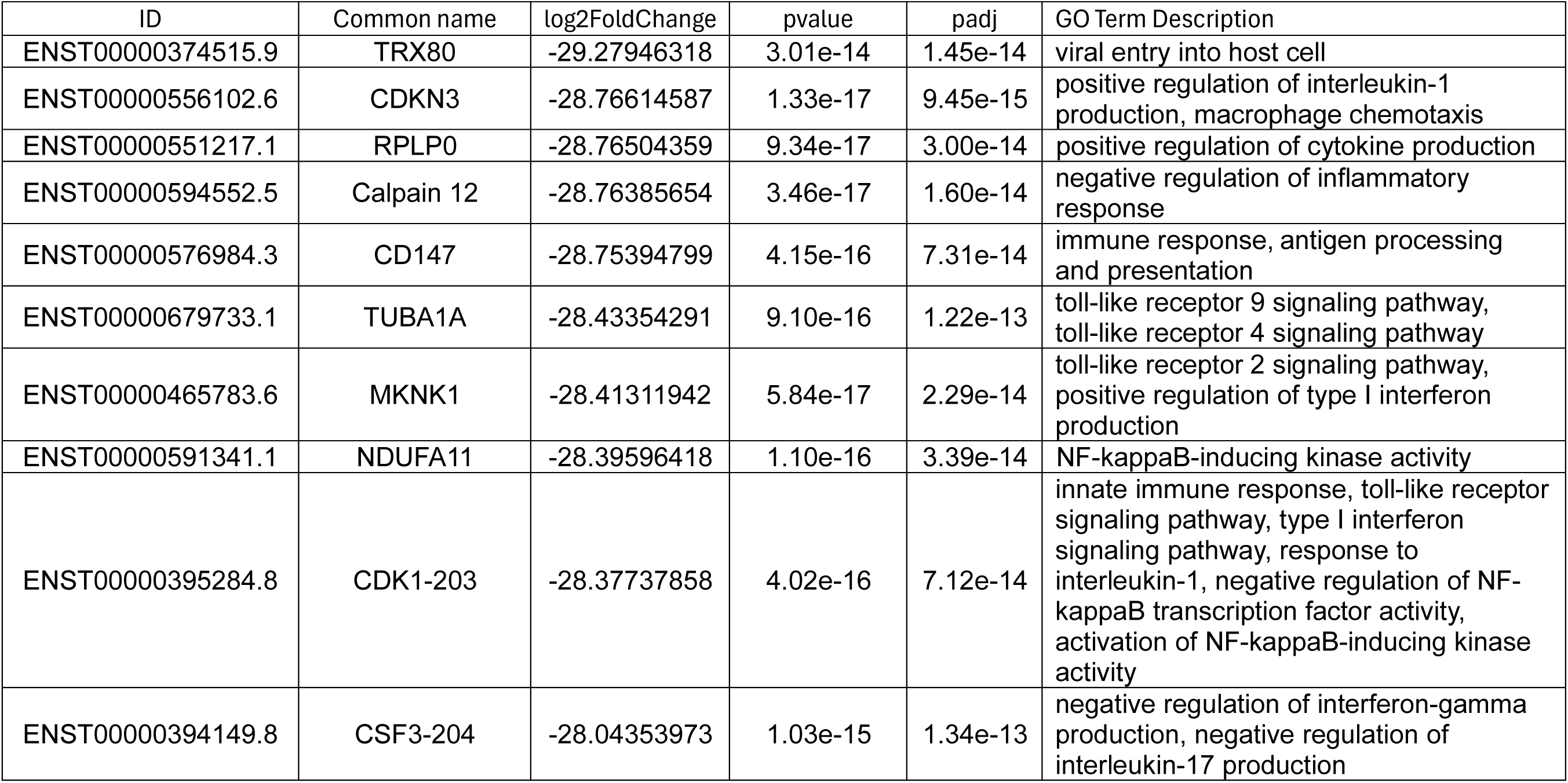
Transcripts Down-Regulated in Response to IAV Exposure and Related to Immune Response and Virus Entry. The transcript ID follow the Ensembl database nomenclature ^71^. The log2 fold change and the adjusted p-value was determined using DESeq ^44^ and the method of Benjamini-Hochberg^70^.

#### Differential Transcript Expression

Table 2 lists the top 10 most significantly upregulated and downregulated transcripts related to immune response and virus entry. The many upregulated transcripts with Gene Ontology (GO) terms related to virus entry and attachment and cytokine signaling indicate that this experiment captured a snapshot of the early stages of exposure. Many of the down-regulated transcripts indicate a reduction in the genes that regulate the immune response. The viral RNA transcripts were not detectable in our experiments, which is consistent with an early stage of infection when the virus is bound to cells, but has not yet internalized or begun viral replication. The moderate MOI and submerged cell culture methods used in this study are conducive to studying the response of exposure to virus and virus binding to the cells, but not viral replication effects *per se*. The importance of cells responding to virus prior to active replicating infection has been highlighted in several recent single cell transcriptome profiles of bystander cells ^45–47^. In low MOI conditions, many cells remain uninfected but still respond to the virus. Interestingly, bystander cells and infected cells show different transcriptomic profiles in influenza infections ^45^. In addition to identifying transcripts expressed in response to exposure to influenza, our approach with nanopore DRS provides information on nucleotide modifications that is not yet technically possible in single cell studies.

#### m6A Methylation Patterns

Figure 2 shows the distribution of m6A methylation in different types of RNA purified from HBECs exposed or not exposed to influenza. Overall, analysis of methylation of protein coding transcripts demonstrates a preference for the GGACA motif as a target for modification with the center A being the nucleotide that is methylated. The protein coding transcripts show a wide variety of motifs that are m6A methylated, while the lncRNA and introns show a more selective usage of targeted motifs. The lncRNA motifs GGACA and AGACU are preferentially modified in HBECs overall, while enhanced modification of the AGACU motif appears to occur in response to influenza exposure. In contrast, the intron RNA show no change in motif usage with virus exposure. Shifts in motif sequence usage from AAC to GAC have been observed in Zika virus infection and were attributed to a change in substrate specificity of the methylation enzymes ^48^. The sequence motif usage is cell specific and idiosyncratic to environmental conditions. The factors determining m6A site selection remain poorly understood ^4^. The m6A readers, writers, and erasers recognize the sequence motif DRACH or RRACH motif where D is A, G, or U; R is A or G; and H is A, C, or U. DRACH motifs occur approximately 1 in every 57 nucleotides, although only 1 in every approximately 1,000 nucleotides is m6A-methylated ^4^. The nucleotide accessibility, local RNA structure, and overall gene architecture contribute to sequence motif usage. In this case of influenza in HBEC cells, the factors influencing the selection of m6A motifs are distinctly different for lncRNA versus protein coding transcripts and change in response to IAV in the lncRNA transcripts. Figure 3 shows an overlay of differentially expressed transcripts and differentially methylated transcripts between diluent-exposed and influenza-exposed HBECs. This analysis focused on a change in the presence or absence of m6A, although methylation can also change sites and stoichiometry. Unexpectedly, the majority of transcripts that show differential m6A do not change transcript abundance. This suggests that mechanisms to regulate transcription might be independent of post-transcriptional mRNA methylation, which can regulate gene expression at the point of protein translation. Only one transcript has reduced m6A and increased transcript abundance. This transcript, *AHNAK2* (ENST00000557457.1), is part of calcium signaling pathways. Only 2 transcripts, *SEMA4B* and *HSPA5,* have reduced m6A and decreased transcript abundance. *SEMA4B* promotes tumor progression in lung cancer proliferation ^49^. Analysis of the m6A Atlas database confirms that *AHNAK2, SEMA4B,* and *HSPA5* have been identified as m6A-methylated transcripts in stimulated A549 cells using traditional Illumina-based methylation identification techniques under different cellular stress conditions^50^. *HSPA5* is part of the heat shock protein 70 family that has a well-studied m6A methylation at nucleotide 103 in the 5’UTR that facilitates 5’cap-independent translation using an internal ribosome entry site ^51^. Heat shock proteins have several roles in influenza infection and help stabilize viral proteins and viral RNA-protein interactions at every step of the viral life cycle ^52^. Interestingly, in our data, the m6A and expression of *HSP5A* both are decreased. Both differential methylation and differential transcription vary during the course of infections. A comparison of differential methylation and differential transcription in *Pseudomonas aeruginosa* bacterial infections, Herpes Simplex Virus-1 (HSV-1) infection, and lipopolysaccharide treatment highlighted the dynamic changes in host m6A methylation over time and in different infection contexts ^12^. In the case of HSV-1, there was also no observed correlation of m6A methylation and differential transcript expression in the host immune response ^12^. Thus, the methylation of RNA transcripts is a highly dynamic mechanism of gene regulation that may display wide variation over time and in different cellular and viral contexts.

**Figure 1:**
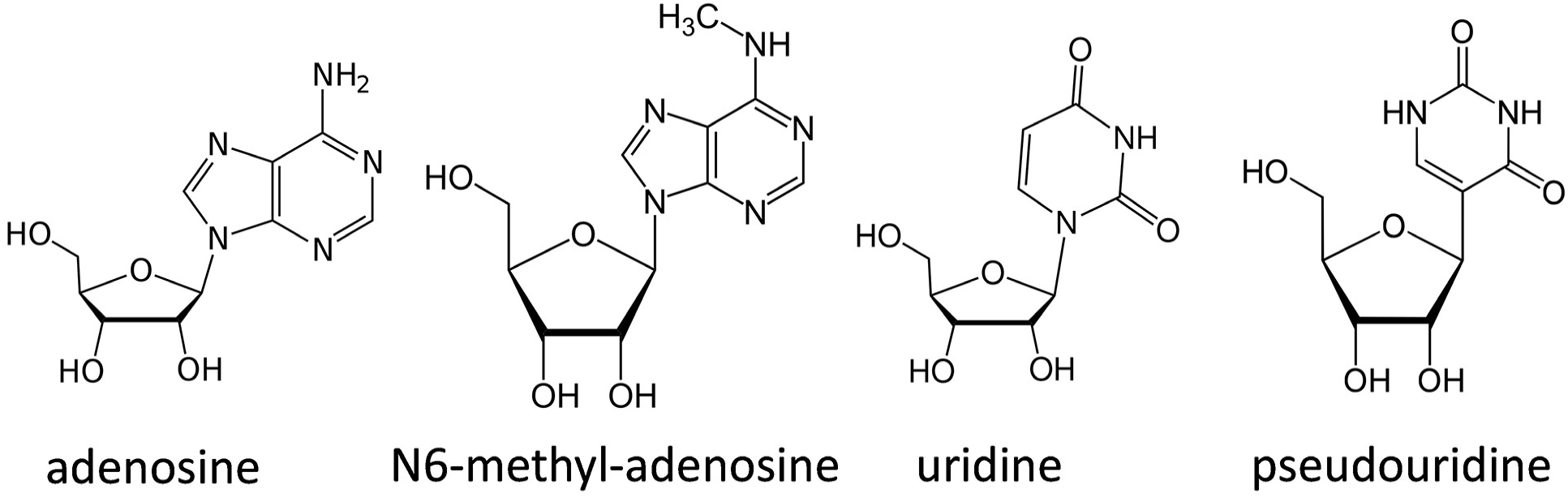
N6-methyl-adenosine (m6A) and Pseudouridine (Ψ) The chemical structure of adenosine, methyl-6-adenosine, uridine, and pseudouridine. N6-methyl-adenosine (m6A) can be directly detected using m6A net analysis of nanopore direct RNA sequence (DRS) data ^25^. NanoSPA identifies both m6A and pseudouridine (Ψ) in DRS data ^27^. In response to exposure to influenza A virus (IAV), both m6A and Ψ change dynamically.

**Figure 2:**
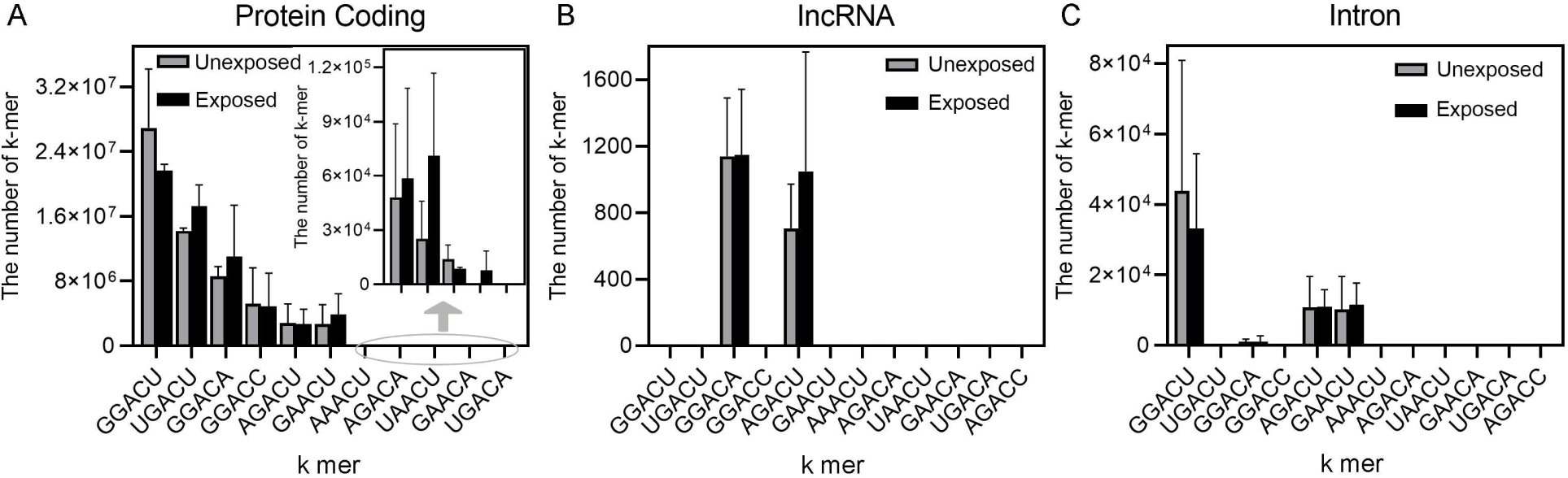
Methylation Motifs in Different Types of RNA after Exposure to IAV. Figure 2. For each type of polyadenylated RNA, a histogram shows the number of each type of sequence motif with the m6A at the center of the motif. GGACU was the most common site of m6A in all types of RNA. The gray bars represent uninfected control samples, and the black bars represent the infected samples.

**Figure 3:**
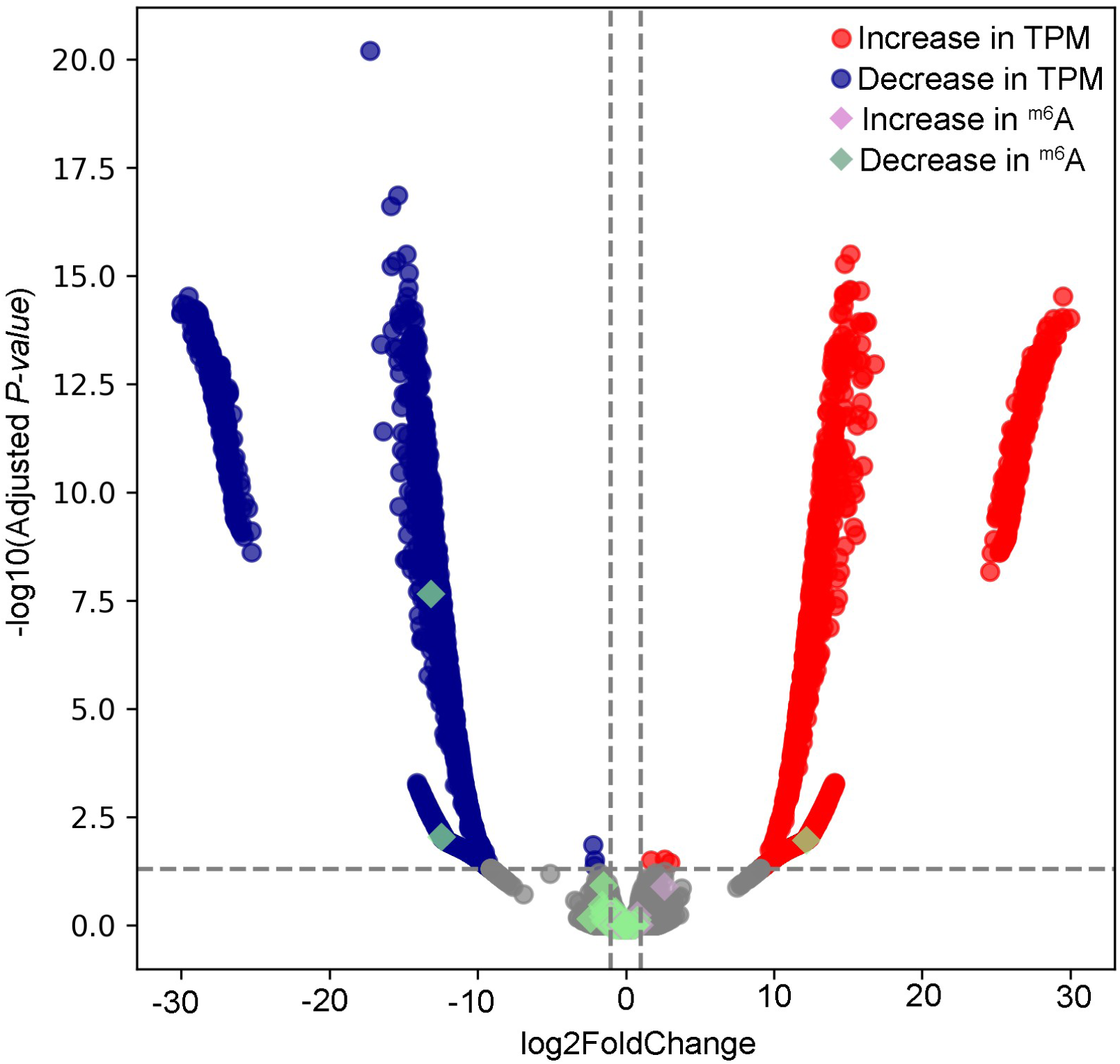
Changes in m6A Methylation are Rare in Transcripts with the Largest Change in Expression in Response to IAV Exposure. Figure 3. Volcano plot shows the log2 fold change versus for log 10 adjusted p-value for differential gene expression in uninfected versus infected HBEC cells. The red circles are overexpressed genes. The blue circles are underexpressed genes. The green diamonds indicate genes that decrease m6A methylation The purple diamonds indicate genes that increase m6A methylation

m6A methylation also does not correlate with alternative RNA splicing in our analysis. Thus, m6A methylation is not likely the main mechanism of regulating the changes in alternative splicing in response to influenza. A similar lack of correlation between m6A methylation and alternative splicing was observed in a study of influenza infection in A549 cells using traditional Illumina sequencing with m6A antibody immunoprecipitation ^53^. The one transcript that is both m6A methylated and changes isoform expression is Calmodulin 2 (*CALM2, ENST0000272298.12)* that has 6 exons and 9 protein isoforms. *CALM2* is part of a system of 3 CALM transcripts that respond to calcium signaling, regulate cell metabolism, and are dynamically regulated by codon usage, mRNA stability, and alternative splicing in response to a wide variety of cellular stresses and disease states^54^. CALM2 was previously identified as an m6A methylated transcript in HEK293T cells^50^. The multiple levels of mRNA regulation for these *CALM* transcripts may enable rapid and robust responses to environmental conditions.

Two lncRNA that regulate immune response are differentially methylated in response to influenza exposure in our RNA nanopore data. Linc01578, also known as *Chaserr*, is methylated at 11 sites in our data. Linc01578 regulates Chromodomain Helicase DNA Binding Protein 2 (*CHD2)* in development and directly binds *NFKB* as part of a positive feedback loop to drive carcinogenesis ^55,56^. Linc00510, also known as *LEADR* and *MIR205HG*, is methylated at 6 sites in our RNA nanopore data. Linc00510 is both a pri-miRNA for miR-205 that regulates the interferon pathway and also a lncRNA that regulates *IRF1* transcription factor activity by binding upstream of the *IRF7* site through *Alu* base pairing ^57,58^. The multiple m6A modifications on these regulatory lncRNA are predicted to change the shape of the lncRNA and disrupt the molecular interactions through which these lncRNA perform regulatory functions.

#### polyA Tail Length

There is an overall decrease in polyA tail length in response to influenza exposure (Figure 4a). Decreased polyA tail lengths are associated with reduced mRNA lifetimes in the cytoplasm and reduced translational efficiency. In our current analysis, 410 genes with GO terms related to immunity and immune regulation show a change in polyA tail length. Some notable immune regulation genes with changes in polyA tail length include *NFKB1* and *NFKB2*; *IRF2, IRF3, IRF6, IRF7*, and *IRF9*; *IL4, IL6*, and *IL32*; *STAT1, STAT2*, and *STAT6*; *TLR2*; *ISG20*; *IFITM1 and IFITM2*; *MAV*; and *OAS1*. Figure 4B shows an overlay of differentially expressed transcripts and differential polyA tail length. Interestingly, the genes with the most change in gene expression are less likely to show a change in polyA tail length. This observation suggests that there are distinct, independent mechanisms of transcriptional and post-transcriptional regulation of host response genes.

**Figure 4:**
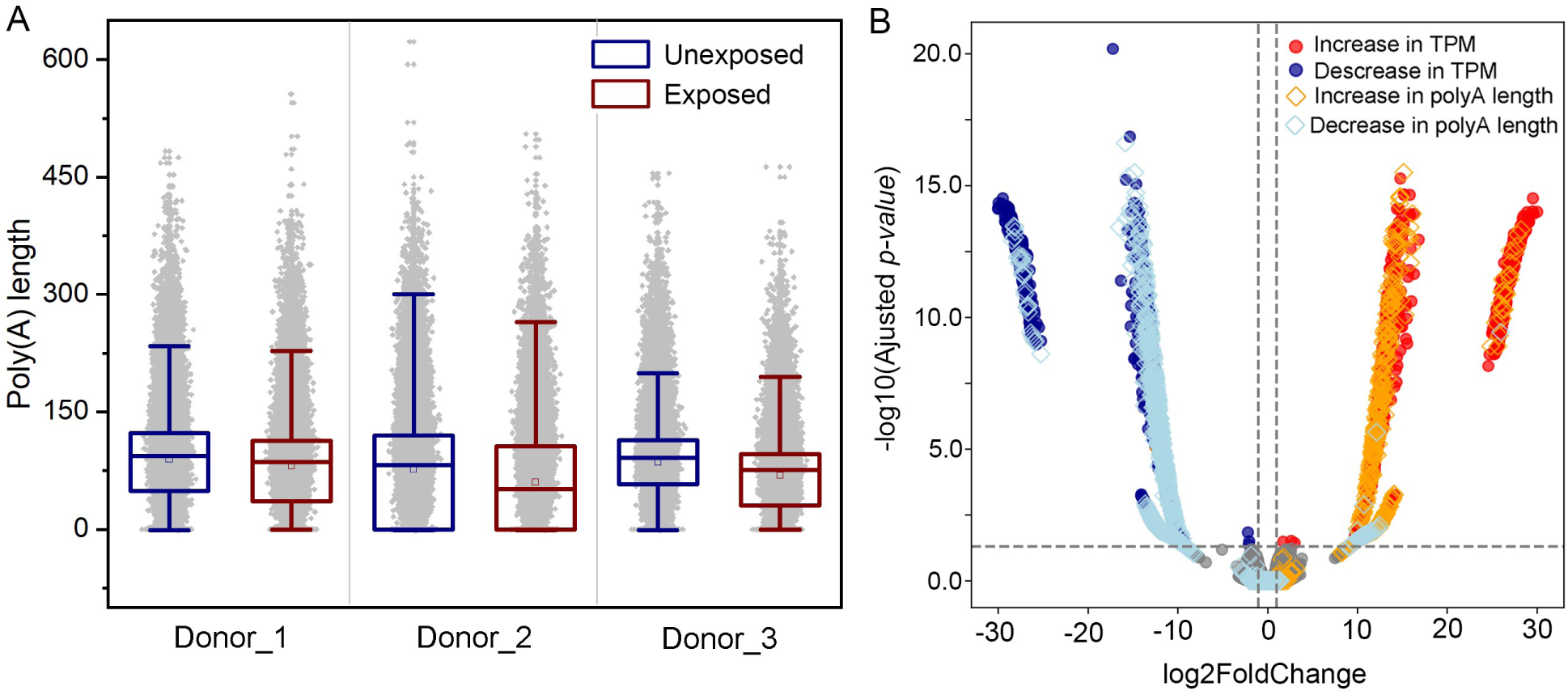
Global Changes in polyA Tail Length with Influenza Exposure. A. Distribution of polyA tail length in each data set. B. Volcano plot shows the log2 fold change versus for log 10 adjusted p-value for differential gene expression in uninfected versus infected HBEC cells. The red circles are overexpressed genes. The blue circles are underexpressed genes. The orange and light blue diamonds indicate transcripts that also change polyA tail length.

#### Alternative Splicing

The longread nanopore data unambiguously identifies the isoforms of transcripts that have multiple alternative splice sites. Among the top 12 differentially expressed transcripts with multiple isoforms (Table 3), the transcript for ISG12 (Interferon Stimulated Gene 12) is significantly upregulated with a 7.83 log2 fold change in response to influenza exposure. ISG12 is known to suppress viral infection through binding to nuclear receptor NR4A1 and preventing nuclear export ^59^. ISG12 has 8 isoforms with complex alternative splicing (Figure 5) ^60^. The long-read sequencing in the direct RNA sequencing greatly facilitates analysis of the changes in expression for each isoform of ISG12.

**Figure 5:**
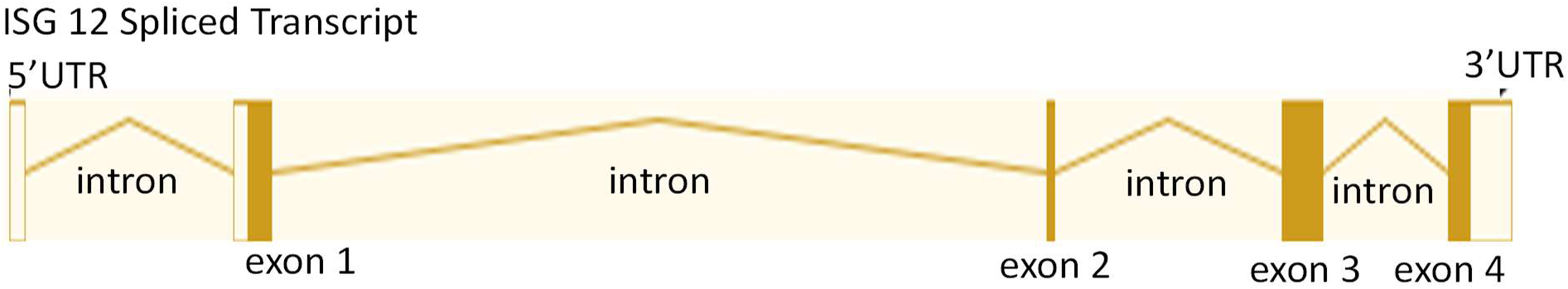
Alternative Splicing of ISG12 (IFI27-213) in Response to IAV Exposure. Transcript ISG12. Interferon Stimulated Gene 12 (ISG12) is also known as IFI27-213 (ENST00000621160.5). Interferon α Inducible Protein 27 is one of 8 isoforms. The DNA sequence spans 5.96 kb. The gene has 4 splice sites. 4 introns, 5 exons, and 4 coding exons (filled boxes). The primary transcript is the longest and has 652 nucleotides that code for a protein with 122 amino acids^60^. The filled boxes indicate the coding exons expressed as a protein in response to IAV, thus highlighting the specific isoform observed to increase expression in our data. Such complex alternative splicing is difficult to identify in short-read sequencing with data from reads 100-200 bp in length. In short-read sequencing, few reads overlap the splice site junction and can be mapped unambiguously to a specific isoform.

**Table 3:** Alternatively Spliced Transcripts Related to Immune Response in Response to IAV Exposure. The transcript ID follow the Ensembl database nomenclature ^71^. The log2 fold change and the adjusted p-value was determined using DESeq ^44^ and the method of Benjamini-Hochberg^70^.

| ID | Common name | log2FoldChange | pvalue | padj | GO Term Description |
| --- | --- | --- | --- | --- | --- |
| ENST00000621160.5 | ISG12 | 7.834827 | 0.000109007 | 0.026674 | immune system process, defense response to virus, type I interferon signaling pathway, innate immune response |
| ENST00000307407.8 | IL8 | -7.54897 | 0.000391257 | 0.059838 | CXC motif chemokine ligand 8 |
| ENST00000393820.2 | Nucleophosmin 1 | 7.999073 | 0.000779847 | 0.082042 | NF-kappaB binding, positive regulation of NF-kappaB transcription factor activity |
| ENST00000495461.6 | SeleneOK | 7.463205 | 0.000966527 | 0.090965 | positive regulation of defense response to virus by host, macrophage derived foam cell differentiation, positive regulation of interleukin-6 production, positive regulation of tumor necrosis factor production, T cell proliferation, T cell migration, positive regulation of neutrophil migration, neutrophil migration, positive regulation of T cell migration |
| ENST00000476133.6 | S100 calcium binding protein A13 | -7.73135 | 0.001269425 | 0.098032 | positive regulation of cytokine production, positive regulation of interleukin-1 alpha production, positive regulation of I-kappaB kinase/NF-kappaB signaling |
| ENST00000525643.7 | IL-32 | -7.33409 | 0.0033791 | 0.168748 | cytokine activity, defense response, immune response |
| ENST00000234875.9 | L22 | -11.2251 | 0.004064586 | 0.181123 | alpha-beta T cell differentiation |
| ENST00000300289.10 | PDIA3 | -7.10807 | 0.004556396 | 0.186878 | cellular response to interleukin-7 |
| ENST00000280357.12 | IL18-201 | -7.09312 | 0.004878905 | 0.195541 | Interleukin 18 |
| ENST00000367351.4 | RAET1H | -6.76689 | 0.009165006 | 0.258238 | positive regulation of T cell mediated cytotoxicity, immune system processes, antigen processing and presentation of endogenous peptide antigen via MHC class Ib, immune response, natural killer cell activation |

### Transcriptional Landscape of Nonpolyadenylated RNA

In response to influenza exposure, expression of nonpolyadenylated noncoding RNA is also changed. We recovered nonpolyadenylated RNA from the polyA purification step, sequenced the transcripts with nanopore sequencing after *in vitro* polyadenylation, and filtered out any protein coding sequences that could be fragments of mRNA transcripts in the bioinformatic analysis. Figure 6 and SI Figure 2A show the distribution of all types of RNA after filtering out any protein-coding RNA transcripts. After the bioinformatic filtering, the distribution of types of RNA is similar in both exposed and unexposed samples. The most common nonpolyadenylated noncoding RNA include 7SL, snRNA, snoRNA, processed pseudogenes, ribozymes, lncRNA, and 7SK.

**Figure 6.**
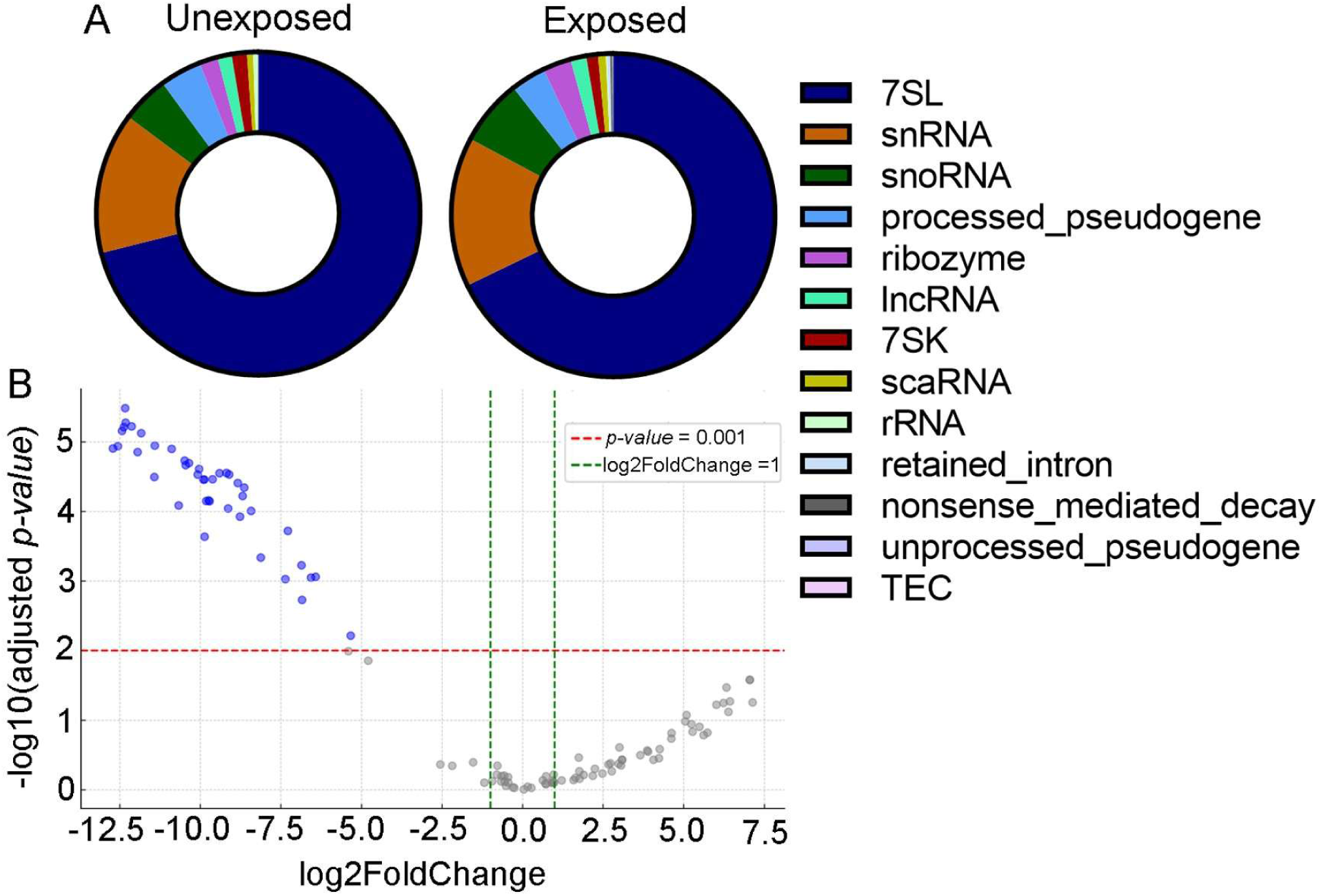
Nonpolyadenylated Noncoding RNA Transcriptome Profiles in Response to IAV Exposure. A. Distribution of types of nonpolyadenylated noncoding RNA in uninfected and PR8-infected HBEC cells by TPM. The total number of unique transcripts is 114. Key: 7SL dark blue; small nuclear RNA (snRNA) orange; small nucleolar RNA (snoRNA) dark green; processed pseudogenes sky blue; ribozymes purple; long noncoding RNA (lncRNA) lime green; 7SK RNA red; small Cajal body associate RNA (scaRNA) gold; ribosomal RNA (rRNA) light green; retained intron light gray; nonsense mediated decay red; unprocessed pseudogenes, light purple; to be experimentally confirmed (TEC), pink. B. The log2 fold change and the adjusted p-value was determined using DESeq ^44^. The dots in blue are statistically significant with a p-adjusted value cutoff of 0.05 using the Benjamini-Hochberg method ^70^.

The most abundant RNA is 7SL RNA. 7SL RNA is a Pol III transcript that normally binds signal recognition particle proteins SRP9 and SRP14. However, unbound 7SL RNA is also a substrate for RIG-I ^61^. In RSV, VSV, and hMPV infections in A549 cells, 7SLRNA is 5mC methylated and activates the RIG-I mediated IFN response ^61^. 7SL RNA can also be m6A methylated in some contexts ^62^, although m6A methylation in our data was not observed with high probability.

Figure 6B shows a volcano plot of the differentially expressed nonpolyadenylated noncoding RNA in response to virus exposure. Interestingly, no transcripts are induced with statistical significance, although many transcripts are downregulated in response to influenza exposure. Table 4 lists the top ten most downregulated nonpolyadenylated noncoding RNA transcripts, which includes snoRNA of the H/ACA type (snoRA2A, snoRA33, snoRA23, snoRA74D), one type of 7SL RNA (rn7SL5P), small nuclear RNA RNU4ATAC, small Cajal body-specific RNA 7 (scaRNA7, a subtype of snoRNA), and 5.8S rRNA. qRTPCR results confirmed a decrease in snoRA23 and snoRA33 in a nonsmoker donor. Dysregulation of snoRNA has been previously reported for PR8 influenza infection in A549 cells ^62^, but no clear mechanism was proposed. In addition to guiding pseudouridylation, snoRNA also are used by influenza viruses in cap snatching ^63–65^. Furthermore, snoRNA binding sites correlate with hotspots for recombination in the hemagglutinin genomic RNA ^66^. Thus, snoRNA have multiple functions in addition to pseudouridylation of ribosomal RNA that play a role in the virus-host interactions in influenza.

**Table 4.** Most Down-Regulated Nonpolyadenylated Noncoding RNA in Response to IAV Exposure. The transcript ID follow the Ensembl database nomenclature ^71^. The log2 fold change and the adjusted p-value was determined using DESeq ^44^ and the method of Benjamini-Hochberg^70^.

| Transcript ID | Common Name | Log2FC | Adj. P-value |
| --- | --- | --- | --- |
| ENST00000383885.1 | SNORA2A | -12.7089 | 0.000144 |
| ENST00000581458.2 | RN7SL5P | -12.5555 | 0.000144 |
| ENST00000363664.1 | SNORA33 | -12.4268 | 0.000142 |
| ENST00000580972.1 | RNU4ATAC | -12.3731 | 0.000142 |
| ENST00000458797.1 | SCARNA7 | -12.3306 | 0.000142 |
| ENST00000365128.1 | SNORA23 | -12.3157 | 0.000142 |
| ENST00000612463.1 | RNA5-8SN2 | -12.1294 | 0.000142 |
| ENST00000619471.1 | RNA5-8SN1 | -11.9426 | 0.000145 |
| ENST00000610460.1 | 5 8S rRNA | -11.8317 | 0.000142 |
| ENST00000516404.1 | SNORA74D | -11.4238 | 0.000172 |

*Pseudouridylation.* Consistent with the downregulation of H/ACA type snoRNA, there is a decrease in pseudouridylation in some nonpolyadenylated ncRNA. Pseudouridylation was identified using the NanoSPA software that detects both m6A and pseudouridylation ^42^. The program accurately identified well-known pseudouridylation sites on 5.8S rRNA at nucleotides 69 and 55 ^67^ with 95% and 85% probabilities, respectively. Thus, the 90% probability cutoff that we used is a conservative identification of pseudouridylation sites. In addition to several ribosomal RNA pseudogenes, two lncRNA were differentially pseudouridylated in response to influenza (SI Table 1). Two lncRNA, linc00273 and novel lncRNA ENST 00000625598.1, are highly pseudouridylated in unexposed cells but not in exposed cells. Linc00273 has 19 high probability pseudouridylated sites in unexposed cells, but none in exposed cells. Linc00273 is transcriptionally induced by STAT3 and plays a role in M2 macrophage activation ^68^. Linc00273 also acts as molecular sponge for miR-200a-3p and promotes lung cancer metastasis^69^. The expression of linc00273 does not change significantly, only its pseudouridylation. This lincRNA is not polyadenylated; and thus, this ncRNA may not have been previously identified as having a role in response to influenza exposure. ENST00000625598.1 is a 923-nucletoide novel antisense lncNRA whose function is yet unidentified. In diluent-exposed cells, the transcript has 16 high probability pseudouridylation sites, while in IAV-exposed cells the lncRNA has no predicted psuedouridylation sites. Further research may reveal the role of these lncRNA and their pseudouridylation in influenza pathogenesis.

#### m6A methylation

Only one nonpolyadenylated lncRNA was predicted to be differently m6A methylated using nanoSPA. Transcript ENST00000629969.1 is a 923 nucleotide antisense lncRNA with no known function. Nucleotide 492 of this transcript is identified as m6A methylated, but not in response to influenza. Interestingly, this modification is predicted to destabilize the RNA secondary structure. Figure 7 shows the predicted centroid RNA secondary structure for the unmodified and modified sequences. The m6A modification reduced the probability of base pairing both in its local helix and at other places in the transcript. One helix adjacent to a large junction is no longer predicted to form at all. This example highlights how m6A methylation could change RNA structure and thereby interactions with small molecule drugs and proteins.

**Figure 7.**
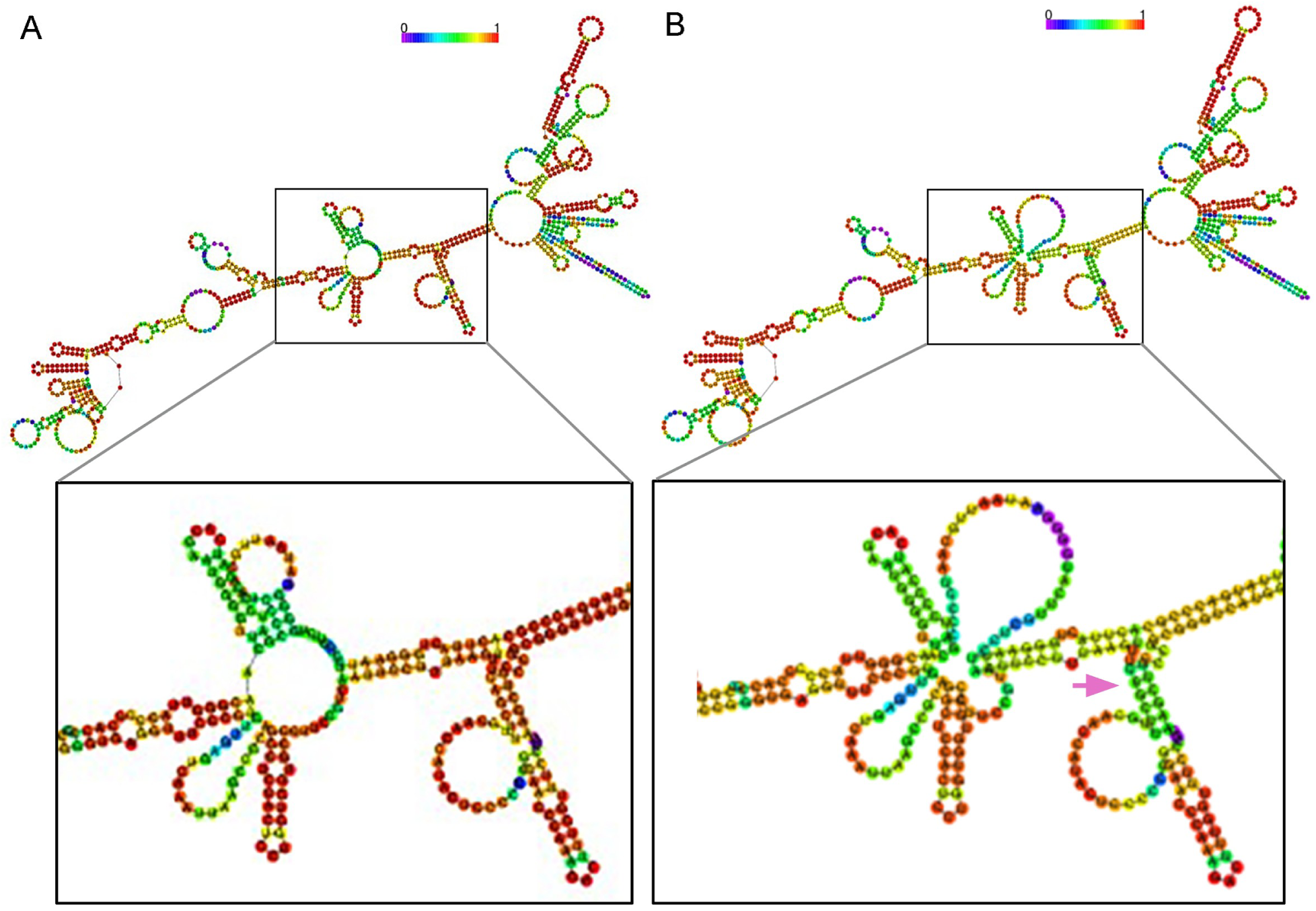
Predicted Structural Effects of m6A in a lncRNA that Responds to IAV Exposure. Predicted centroid secondary structures for ENST00000629969.1 without (A) and with (B) m6A at nucleotide 492. The color of the nucleotide indicates the probability of the nucleotide being in a base pair or unpaired with red being a probability of 1 and purple being a probability of zero. Structures were generated using Vienna rnafold with default parameters in May 8,2024 ^72^. The pink arrow in B indicates the location of the m6A modification. The Vienna program uses the thermodynamic parameters from ^73^.

## Conclusion

This study profiles the epitranscriptomic changes in human primary bronchial epithelial cells in the early stages of influenza virus exposure. Notably, all the changes in mRNA transcripts, polyadenylated noncoding RNA, and nonpolyadenylated noncoding RNA, observed in this study required only exposure and not active virus replication. The epitranscriptomic modification of noncoding RNA extends the mechanistic repertoire of noncoding RNA gene regulation to include changing RNA interactions at the post-transcription stage in response to environmental cues, stresses, and infections. There is a decrease in H/ACA type snoRNA and a decrease in pseudouridylation of several noncoding RNA. The m6A methylation patterns are distinctly different for coding and noncoding RNA. Unexpectedly, m6A methylation did not correlate with alternative splicing, polyA tail length, or differential expression. m6A modification does have important effects on the transcriptome, as the m6A and pseudouridylation modifications may change RNA cellular stability, RNA folding, or RNA interactions with proteins, small molecules, and other RNA. The potential for m6A to alter RNA secondary structure is demonstrated for a novel lncRNA. The use of direct RNA nanopore sequencing highlights the ability to identify in one experiment the differential expression, modifications, polyA tail length, and unambiguous splicing isoform in the human transcriptome and directly test correlations of modifications and the mechanisms with which modification fine tune gene expression. The ability to identify multiple modifications on a long read of an individual RNA molecule surpasses other techniques based on cleaving or fragmenting the nucleic acids for short read sequencing. For example, linc00273 has multiple pseudouridylation sites, and this example provides a foundation for further testing the functional role of this extensive pseudouridylation. As a single molecule experiment, nanopore direct RNA sequencing has the potential to reveal individual variations that bulk sequencing would obscure. This study presents an epitranscriptome profile that provides a foundation for future discovery of the mechanisms through which noncoding RNA and RNA modifications fine-tune the host response to influenza infections and exposure.

## Acknowledgement

Research reported in this publication was supported by the National Institute of General Medical Sciences of the National Institutes of Health under Award Number P20GM103648 and P30GM149368. The content is solely the responsibility of the authors and does not necessarily represent the official views of the National Institutes of Health. Research reported in this publication was supported by the Merit Review Program of the Department of Veterans Affairs, grant number I01 BX005023 (JPM); Clinical Innovator Award from the Flight Attendant Medical Research Institute # 113052 (JPM); resources and assistance from the Civil Aerospace Medical Institute, Federal Aviation Administration (JPM); a Seed Grant Award from the Presbyterian Health Foundation (JPM and SJS), the Dodge Family College of Arts and Sciences Collaborative Fellowships Award at the University of Oklahoma (SJS); and an MCI supplement to NSF TRTech Award #2023310 (SJS). The funders had no role in study design, data collection and interpretation, or the decision to submit the work for publication. We thank the Consolidated Core Lab at the University of Oklahoma for the use of the facility that provided nanopore sequencing resources and equipment.

## Author Contributions

DW and SJS wrote the main manuscript text. DW, ML, and SJS prepared figures. All authors reviewed the manuscript.

## Data Availability

The data has been deposited to the European Nucleic Acid Archive PRJNA1134237.

