## Supporting Information for "Nanopore Direct RNA Sequencing Reveals Virus-Induced Changes in the Transcriptional Landscape in Human Bronchial Epithelial Cells"

Running Head: Nanopore RNA Sequencing Transcriptional Landscape

Key words: Direct RNA nanopore sequencing, m6A methylation, virus-host interaction, influenza  in primary human bronchial epithelial cells, epitranscriptomics

**SI Table 1: Nonpolyadenylated Noncoding RNA with Changes in Pseudouridylation Induced by IAV Exposure**

| Transcript | Common name | Ψ sites, uninfected | Ψ sites, infected |
|---|---|---|---|
| ENST00000539813.1 | Linc00273 | 323, 350, 430, 553, 583, 674, 700, 716, 758, 773, 833, 838, 851, 876, 903, 917, 969, 1033, 1042 | none |
| ENST00000625598.1 | Novel lncRNA | 45, 46, 61, 68, 291, 292, 423, 478, 484, 494, 593, 613, 692, 871 | none |
| ENST00000445125.2 | 18 S ribosomal pseudogene Y chromosome | 35, 99, 109, 461, 517, 568, 572, 590, 648, 707, 957, 1108, 1226, 1236, 1339, 1566 | none |
| ENST00000445125.2 | 18 S ribosomal pseudogene Y chromosome | none | 1782 |

*SI Table 1: Pseudouridylation sites were identified using nanoSPA [1] and a cutoff of 90% probability. Only sites consistently identified in both donors are included.*

**SI Table 2: Primers for qRTPCR**

| name | sequence | target |
|---|---|---|
| snoRA23F | 5’TGGTAGCAGTGTCTGTCTGTG | snoRA23 |
| snoRA23R | 5’CCAAGTTACTCTTTGGCCGC | |
| snoRA33F | 5’AAGCCAGCCAATGAATCTGC | snoRA33 |
| snoRA33R | 5’TTGTTATAGCCATTCTCAGGG | |
| BactinF | 5’GCCGGGACCTGACTGACTAC | β-actin |
| BactinR | 5’TTCTCCTTAATGTCACGCACGAT | |

*SI Table 2: Primers for snoRA23 are from reference. Primers for snoRA33 are from reference[2]. Primers for β-actin are from reference[3].*

### SI Figure 1: RNA Quality and Basecalling Accuracy

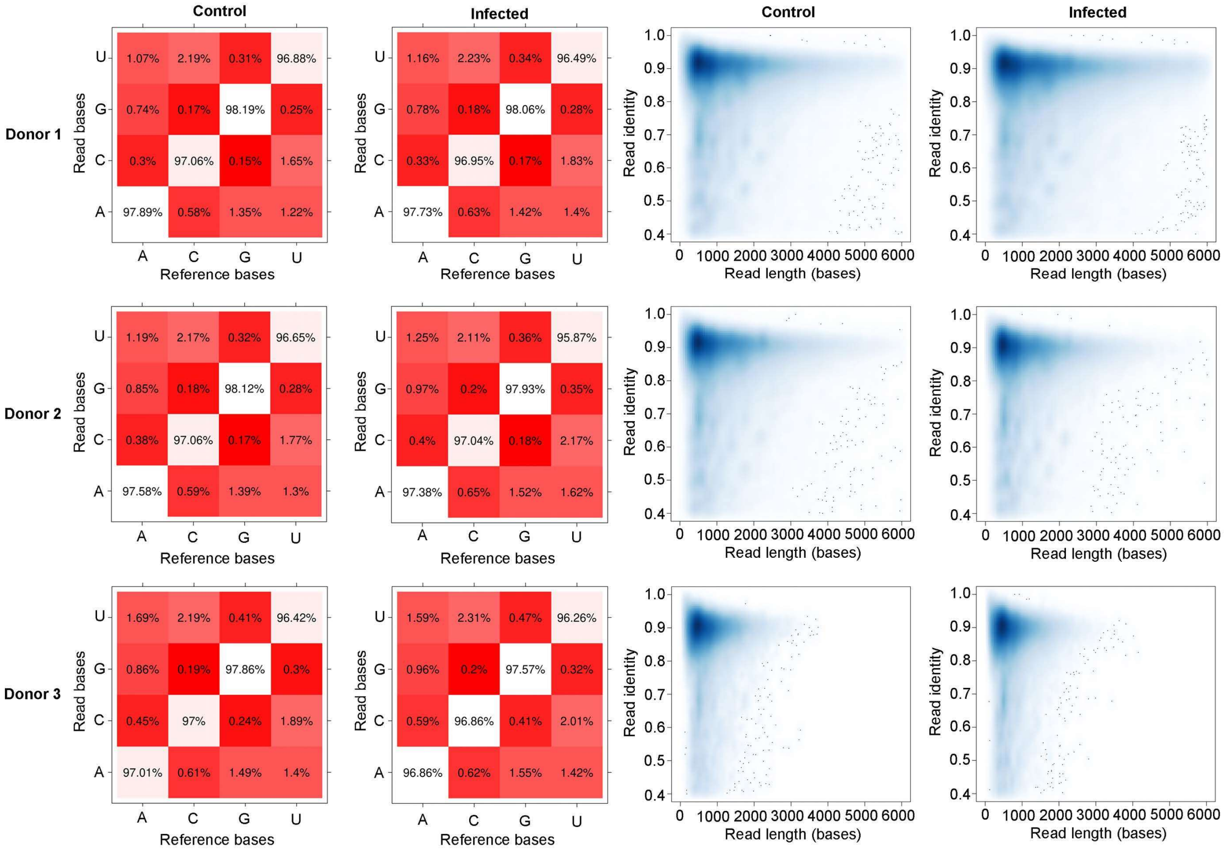


*SI Figure 1: Base calling accuracy plots and normalized read identity versus length plots were calculated following the analysis used by Workman et al.[4].*

**SI Figure 2. Noncoding RNA Profiles Induced by IAV Exposure for Each Donor**

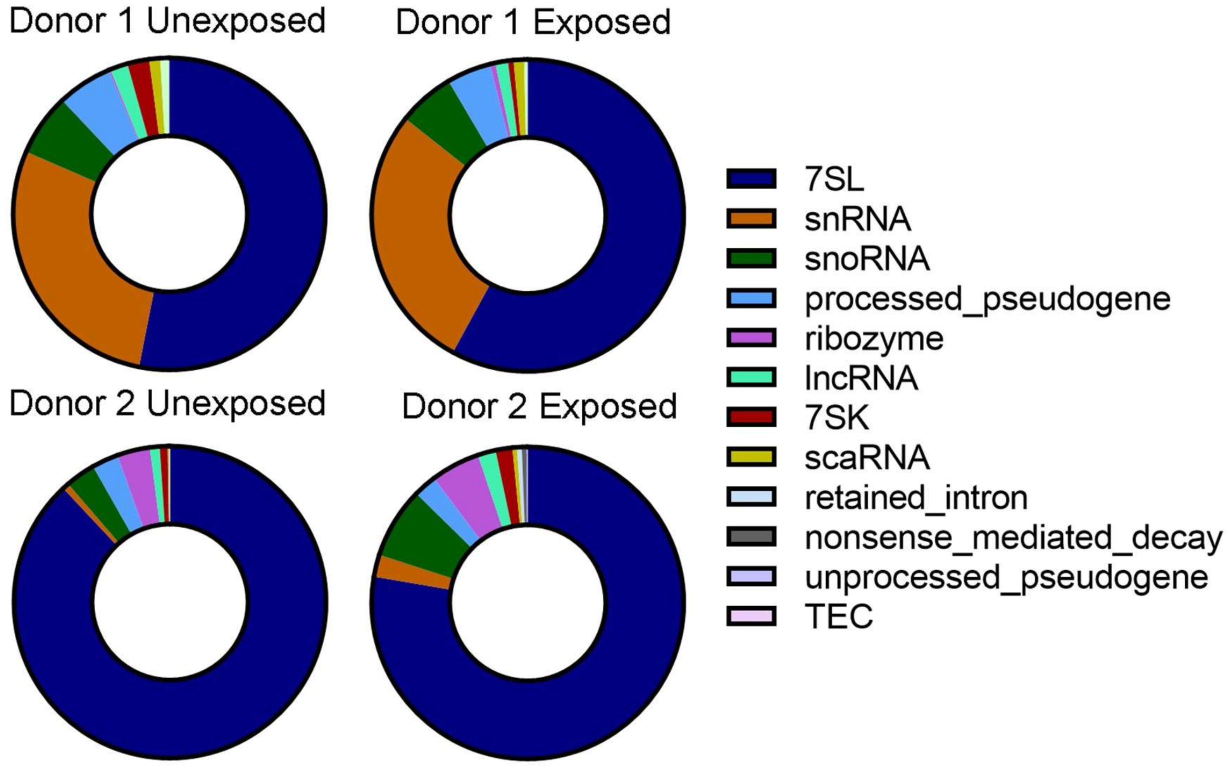


*SI Figure 2. Distribution of types of nonpolyadenylated noncoding RNA in uninfected and IAV-exposed HBEC cells for each donor data set by TPM. Key: 7SL dark blue; small nuclear RNA (snRNA) orange; small nucleolar RNA (snoRNA) dark green; processed pseudogenes sky blue; ribozymes purple; long noncoding RNA (lncRNA) lime green; 7SK RNA red; small Cajal body associate RNA (scaRNA) gold; ribosomal RNA (rRNA) light green; retained intron light gray; nonsense mediated decay red; unprocessed pseudogenes dark gray; unprocessed pseudogenes, light purple; to be experimentally confirmed (TEC), pink.*
